# Abundance and localization of human UBE3A protein isoforms

**DOI:** 10.1101/2020.03.30.016857

**Authors:** Carissa L. Sirois, Judy E. Bloom, James J. Fink, Dea Gorka, Steffen Keller, Noelle D. Germain, Eric S. Levine, Stormy J. Chamberlain

## Abstract

Loss of *UBE3A expression,* a gene regulated by genomic imprinting, causes Angelman Syndrome (AS), a rare neurodevelopmental disorder. The *UBE3A* gene encodes an E3 ubiquitin ligase with three known protein isoforms in humans. Studies in mouse suggest that the human isoforms may have differences in localization and neuronal function. A recent case study reported mild AS phenotypes in individuals lacking one specific isoform. Here we have used CRISPR/Cas9 to generate isogenic human embryonic stem cells (hESCs) that lack the individual protein isoforms. We demonstrate that isoform 1 accounts for the majority of UBE3A protein in hESCs and neurons. We also show that UBE3A predominantly localizes to the cytoplasm in both wild type and isoform-null cells. Finally, we show that neurons lacking isoform 1 display a less severe electrophysiological AS phenotype.

---

Angelman Syndrome (AS) is a rare neurodevelopmental disorder that affects 1 in 15,000 individuals, and is characterized by severe seizures, intellectual disability, absent speech, ataxia, and happy affect^1^. AS is caused by loss of function from the maternally-inherited copy of *UBE3A. UBE3A* is regulated by tissue-specific genomic imprinting - it is expressed exclusively from the maternal allele in neurons and is expressed biallelically in other cell types^2,3^. *UBE3A* encodes an E3 ubiquitin ligase that forms polyubiquitin chains to substrates, targeting them for degradation by the 26S proteasome^4^. In humans, there are three known UBE3A protein isoforms^5^, all of which include the Homologous to E6AP Carboxy Terminus (HECT) domain and are thus capable of functioning as an E3 ligase. Human isoforms 2 and 3 only differ from human isoform 1 by 23 and 20 amino acids, respectively, at their N termini^6^ (Fig 1a). While there has been little published research examining the human protein isoforms in human cells, human isoforms ectopically expressed in mouse indicate that the isoforms likely have differences in localization and function^7–9^. Furthermore, it is currently unknown whether the different human isoforms have different abundances or functions. This knowledge is important as some therapeutic avenues currently being explored for AS involve delivery and expression of exogenous *UBE3A* transgenes using vector-based delivery^10^. For these approaches, knowledge of which of the protein isoforms need to be replaced in AS is essential.

**Figure 1.**
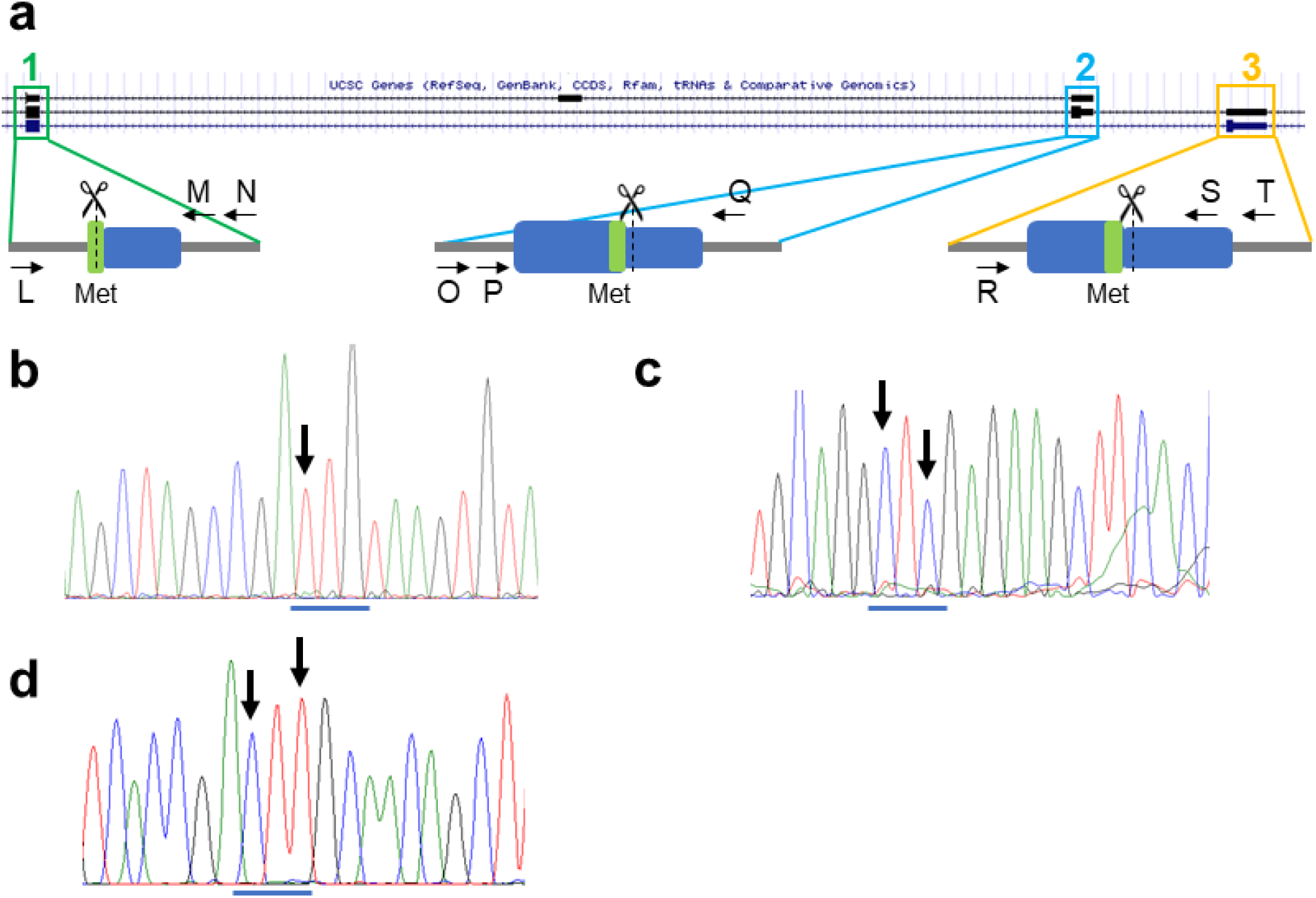
Generation of isogenic isoform-null hESC lines. **a** Schematic illustrating isoform translational start sites and proposed genome editing. Arrows indicate primers used for screening. Scissors indicate CRISPR cut sites. Blue box = exon; grey line = intron; green = methionine used as start site **b-d** Sanger sequencing showing mutation of translational start site (ATG) for isoform 1 (ATG→TTG; **b**), isoform 2 (ATG→CTC; **c**), and isoform 3 (ATG→CTT; **c**). Arrows indicate locations of changed nucleotides. The original ATG start site is underlined in blue. The edited nucleotides changed the original methionine to a leucine.

Recently, Sadhwani and colleagues published three cases of AS in two different families caused by missense mutations at the isoform 1 translational start site^11^. Interestingly, these patients present with milder phenotypes than AS patients with 15q11-q13 deletions or other *UBE3A* loss of function mutations, including normal gait and use of syntactic speech. This data, coupled with the fact that the human and mouse isoforms are not entirely conserved, illustrates the need to study the human UBE3A protein isoforms specifically, as they may play a role in both normal neuronal function and in disease. Here, we have used human embryonic stem cells (hESCs) and their neuronal derivatives to examine the abundance and localization of the three human isoforms. We have also examined whether neurons lacking individual protein isoforms recapitulate any AS phenotypes previously seen in induced pluripotent stem cell (iPSC)-derived neurons. We show that in both hESCs and hESC-derived neurons, all three UBE3A protein isoforms are predominantly localized to the cytoplasm, with low levels of expression in other cellular compartments. We also show that protein isoform 1 is the predominant protein isoform in human cells. Finally, we show that neurons lacking isoform 1 show some, but not all, of the phenotypes displayed by AS iPSC-derived neurons, while loss of isoform 2 or 3 does not produce any phenotypic changes. These results not only provide useful insight into the human UBE3A isoforms, but also provide information important for the development of potential AS therapies.

## Results

### Generation of isogenic hESC lines lacking individual protein isoforms

To study the abundance and localization of the UBE3A protein isoforms, we first generated isogenic hESC lines lacking the individual protein isoforms. Isogenic hESCs and neurons would allow us to minimize molecular and phenotypic differences caused by normal human genetic variation. All three human protein isoforms are full length versions of UBE3A: isoforms 2 and 3 have an additional 23 and 20 amino acids, respectively, at their N terminus. Because of this, the RNAs encoding the isoforms are predicted to be nearly identical aside from the exons that encode for these additional amino acids^6^. The three UBE3A protein isoforms are, however, translated from unique translational start sites^12^. Using CRISPR/Cas9 and singlestranded olignonucleotide (ssODN) templates, we mutated each of the three translational start sites, changing them from methionines to leucines, to prevent the translation of the isoform of interest (**Figure 1**). Because the methionine that serves as the isoform 1 translational start site is present in the other two protein isoforms, we used multiple protein prediction softwares^13,14^ to find an amino acid substitute (leucine) that was predicted to have a benign effect on the other two isoforms (**Supplemental Figure 1**). Indeed, all three isoform-null hESC lines made normal levels of UBE3A mRNA (**Supplemental Figure 2b**).

### Isoform 1 is the most abundant UBE3A protein isoform

To determine the relative abundance of the human UBE3A protein isoforms in hESCs and hESC-derived neurons, we examined total UBE3A protein levels in whole cell lysates prepared from the isoform knockout and isogenic control lines. Our data indicate that isoform 1 is the predominant isoform in both cell types - loss of this isoform produced a significant reduction (84-88%) in total UBE3A levels in whole cell lysates prepared from both hESCs (**Figure 2a,b**) and neurons (**Figure 3a,b**). Loss of isoform 2 or isoform 3, however, did not produce any significant changes in total UBE3A levels. These data indicate that isoform 1 is the most abundant of the three human protein isoforms in hESCs and hESC-derived neurons.

**Figure 2.**
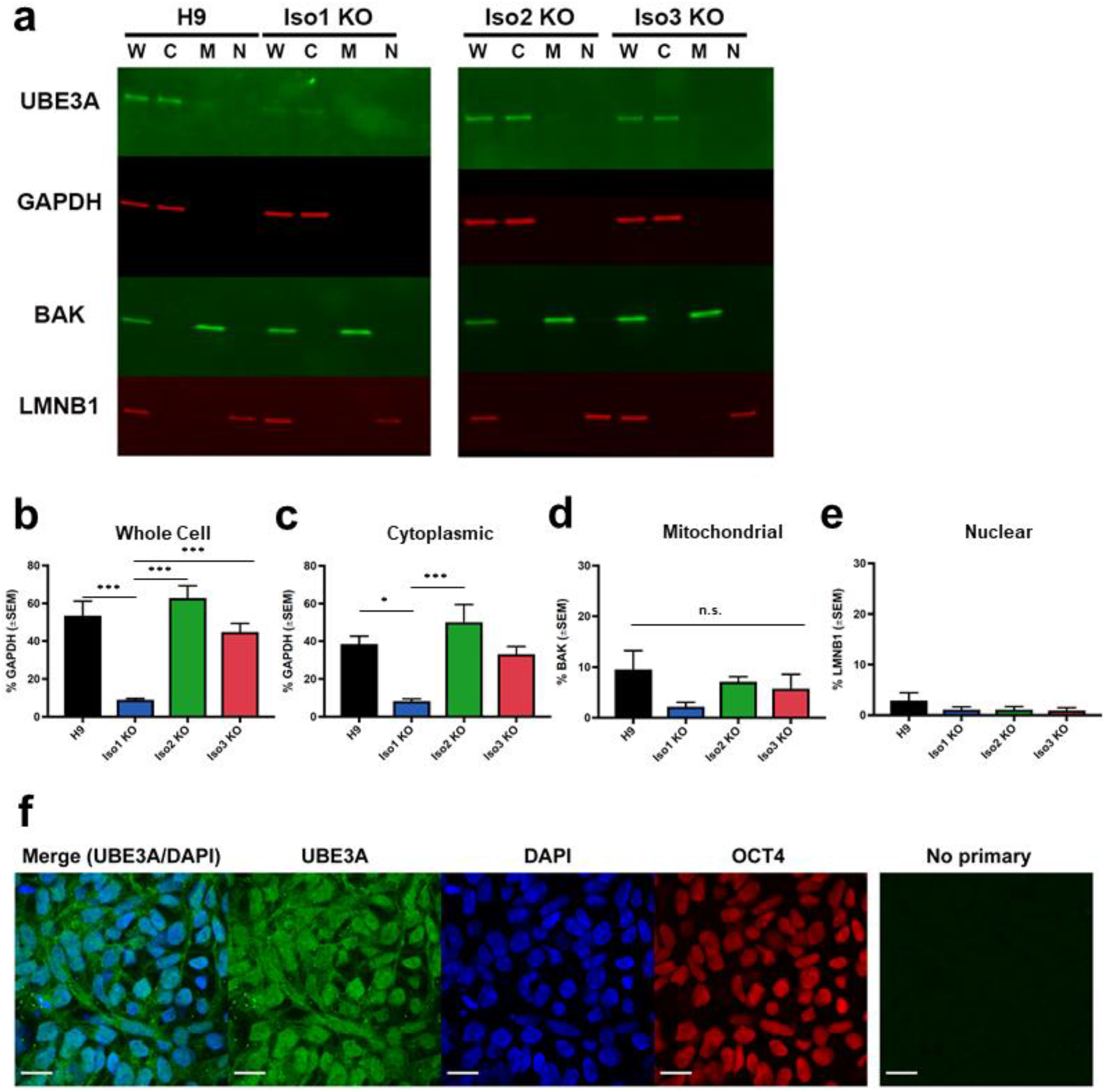
Abundance and localization of UBE3A isoforms in hESCs. **a** Western blot showing UBE3A levels in subcellular fractions from H9 and isoform-null hESC lines. W = whole cell lysate, C = cytoplasmic fraction, M = mitochondrial fraction, N = nuclear fraction. **b-e** Quantification of Western blots for each fraction. Graphs show average percentage of appropriate loading control (n = 4 fractionations). Error bars: standard error of the mean. * p < 0.05 *** p < 0.005 (univariate ANOVA) **f** Immunocytochemistry for UBE3A in H9 hESCS shows that UBE3A is cytoplasmic, in agreement with above fractionation results. Scale bar 20 μm.

### Subcellular localization of the three UBE3A isoforms

Because studies of the mouse protein isoforms indicate differences in isoform localization^7,9^, we sought to determine whether the human protein isoforms also localized differentially. We examined UBE3A abundance in cytoplasmic, mitochondrial, and nuclear fractions in both hESCs and neurons. In both cell types, the bulk of UBE3A protein is found in the cytoplasm, both in normal and isoform-null lines (**Figure 2a-e, Figure 3a-e**). Consistent with our whole cell results, loss of isoform 1 produced a significant reduction of UBE3A from the cytoplasmic fraction (**Figure 2c, Figure 3c**), but not the mitochondrial or nuclear fractions (**Figure 2d/e, Figure 3 d/e**). Loss of isoform 2 or isoform 3 did not cause a significant change in the abundance of UBE3A protein in any cellular fractions when compared to the H9 parent line (**Figure 2a-e, Figure 3a-e**). Since UBE3A in the nuclear fractions was difficult to see, we ran a Western blot using a larger concentration of lysates from wildtype and isoform-null hESCs and neurons that was intentionally exposed to the point of saturation (3 min; **Supplemental Figure 5b**). This showed a faint signal, confirming the presence of UBE3A protein in the nucleus.

**Figure 3.**
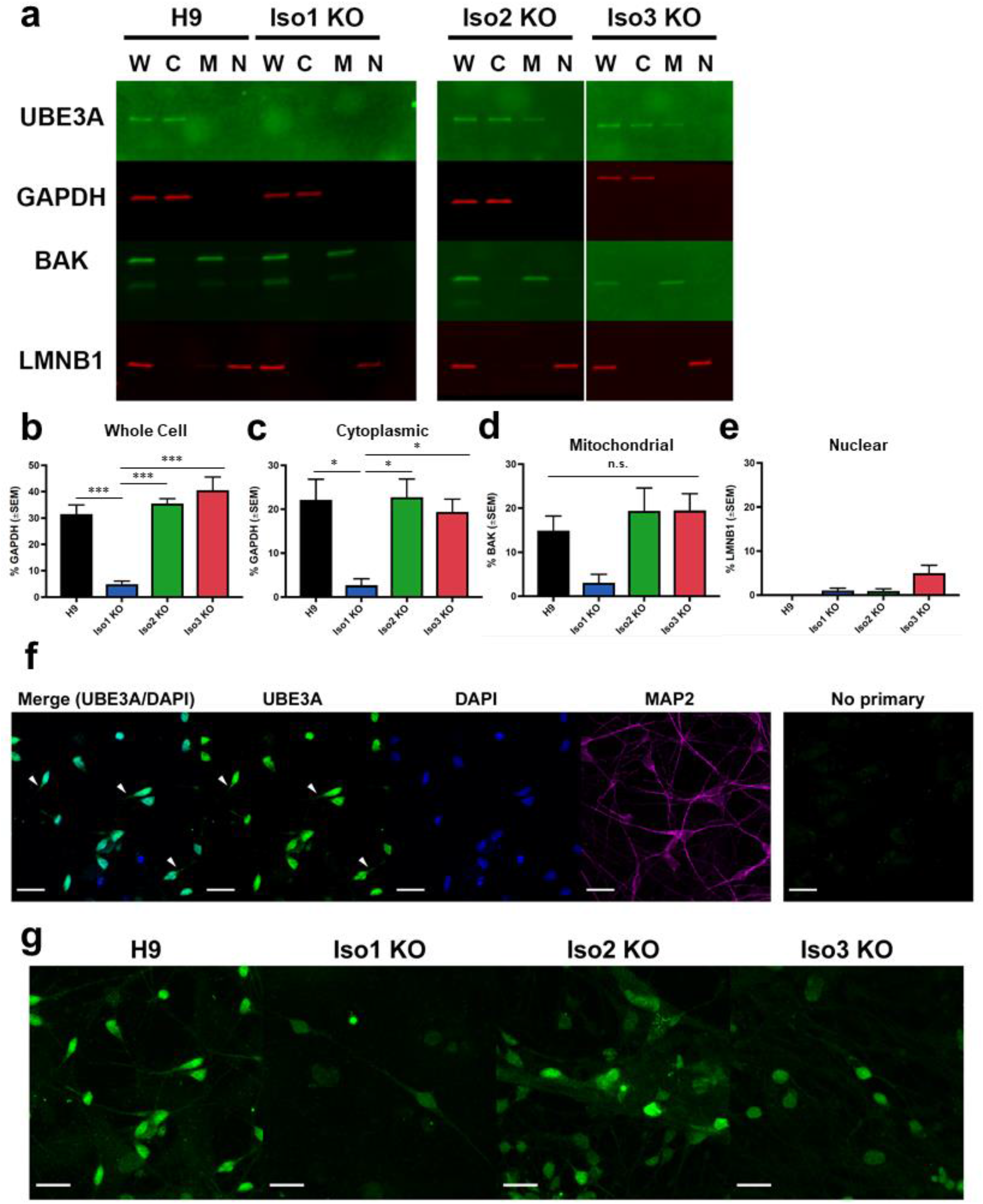
Abundance and localization of UBE3A isoforms in neurons. **a** Western blot showing UBE3A levels in subcellular fractions from H9 and isoform-null neurons. W = whole cell lysate, C = cytoplasmic fraction, M = mitochondrial fraction, N = nuclear fraction. **b-e** Quantification of Western blots for each fraction. Graphs show average percentage of appropriate loading control (n = 3 fractionations). Error bars: standard error of the mean. * p < 0.05 *** p < 0.005 (univariate ANOVA) **f-g** Immunocytochemistry for UBE3A in H9 neurons shows that UBE3A is highly concentrated in the nucleus but also is expressed in the cytoplasm. **f** single plane image, **g** maximum projections of confocal z-stacks. Scale bar 20 μm. Arrowheads indicate areas in the neuron where UBE3A is expressed outside of the nucleus.

We confirmed the fractionation results using wildtype hESCs and neurons along with a second primary antibody against UBE3A, which reacts to a different portion of the protein (N terminus versus HECT domain; **Supplemental Figure 5a**). A third primary antibody (the same antibody used for immunocytochemistry, see below) similarly showed an almost undetectable nuclear band in both wildtype and Isoform1-null hESCs (**Supplemental Figure 5c**) as well as in wildtype neurons (**Supplemental Figure 5d**). Fractionation was also carried out on hESCs harboring a homozygous insertion of a hemagglutinin (HA)-tag into the isoform 1 translational start site of UBE3A. Western blot performed using an anti-HA antibody further confirmed our previous observations (**Supplemental figure 7b**).

We then examined UBE3A localization via an orthologous method: immunocytochemistry. We first stained hESCs for UBE3A and OCT3/4, a transcription factor and pluripotency marker that localizes to the nucleus (**Figure 2f, Supplemental Figure 3**). We included an isogenic UBE3A KO hESC line to serve as a negative control (**Supplemental Figure 2c-f**). Expression of UBE3A in hESCs appears to be distributed across the cytoplasm and nucleus in both normal and isoform-null hESCs, as there is expression of UBE3A inside and outside of the area positive for OCT3/4 staining. We next stained hESC-derived neurons for UBE3A and MAP2, a microtubule protein that localizes to the cytoplasm and is a marker for post-mitotic neurons (**Figure 3f, Supplemental Figure 4a**). Although the strongest UBE3A signal appears in the nucleus, there is clear UBE3A signal in the neuronal processes and in the area of the soma outside of the nucleus (**Supplemental Figure 4a)**. This is especially apparent in maximum intensity projections of confocal z-stacks from normal and isoform-null neurons (**Figure 3g**).

Immunocytochemistry was repeated in neurons derived from wildtype and isoform-null hESCs using a different primary antibody against UBE3A. These results (**Supplemental Figure 6a**) confirmed our previous observations: while there was strong UBE3A signal in the nucleus, UBE3A can also be seen outside of the nucleus in both the soma and processes. Wildtype neurons co-labeled with antibodies against UBE3A and NeuN, which is expressed predominantly in the nucleus in mature neurons, showed intense nuclear UBE3A signal, as evidenced by co-localization with NeuN, but also revealed UBE3A signal in the soma and neuronal processes (**Supplemental Figure 6b**). Immunocytochemistry performed using anti-HA antibodies in neurons derived from UBE3A-HA-tagged hESCs similarly labeled the nucleus, soma, and neuronal processes (**Supplemental Figure 7c**).

### UBE3A is diffusely localized throughout nuclei and cytoplasm in stem cells and neurons

To make sense of seemingly disparate UBE3A localization results, we examined UBE3A localization in neurons by transmission electron microscopy (TEM). These results showed diffuse UBE3A puncta in the soma both within and outside of nucleus, as well as in the neuronal processes (**Supplemental Figure 8**).

### Isoform 1-null neurons display mild AS electrical phenotypes

Previously we established that AS iPSC-derived neurons exhibit a phenotype of impaired physiological maturation, as indicated by a more depolarized resting membrane potential (RMP), immature patterns of action potential firing, and decreased frequency of spontaneous excitatory synaptic activity^15^. We last wanted to determine whether human neurons lacking the individual UBE3A protein isoforms displayed any of the same AS phenotypes. Based on our results examining the abundance of the isoforms (**Figure 3a-b**), and our knowledge of the AS patients with isoform 1 translational start site mutations^11^, we hypothesized that loss of isoform 1 would produce AS phenotypes in hESC-derived neurons while loss of the other two isoforms would likely would not produce any phenotype. We examined these three phenotypes in our isoform-null and isogenic control neurons at 12 weeks of differentiation. Isoform 1 KO neurons displayed a more depolarized resting membrane potential compared to controls, while there were no differences in the isoform 2 KO or isoform 3 KO neurons (**Figure 4a**). Interestingly, there were no statistically significant differences in action potential firing (**Figure 4b**) or synaptic activity (**Figure 4c**) in the any of the isoform-null neurons, indicating that the isoform 1 KO neurons only display limited AS electrical phenotypes.

**Figure 4.**
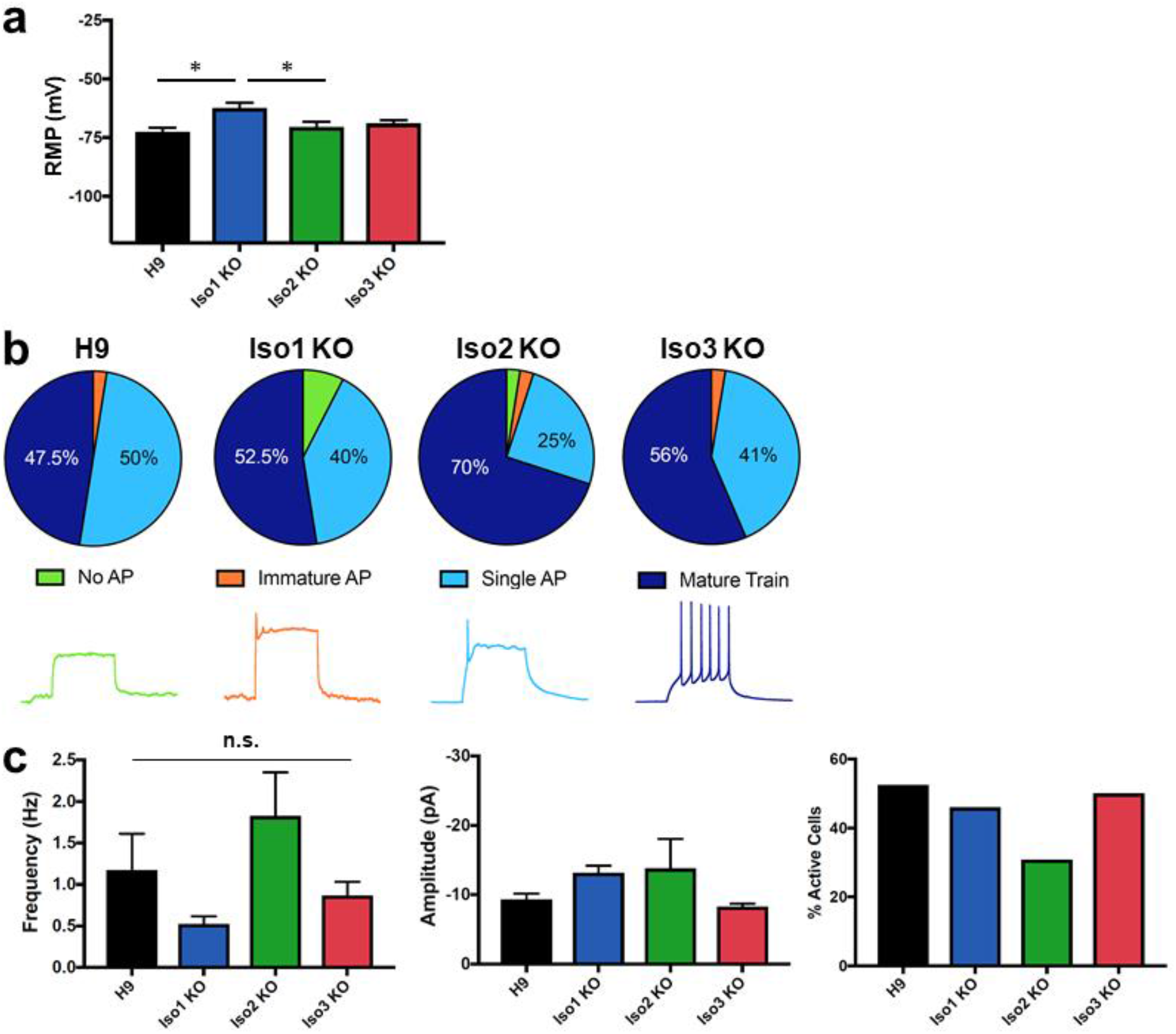
Isoform 1-null neurons partially recapitulate AS cellular phenotypes. **a** Isoform-1 null neurons have a more depolarized resting membrane potential. * p < 0.05 (univariate ANOVA) **b** No significant differences in action potential (AP) firing upon loss of any UBE3A isoforms, Top: AP characterization in isoform-null neurons; Bottom: sample AP traces **c** Isoform-null neurons do not have any significant differences in spontaneous excitatory synaptic activity (left, middle). n = 39 (Iso3 KO) or 40 (all other lines) cells on 3 coverslips per cell line; only active neurons were included in these analyses (right). All error bars: standard error of the mean.

## Discussion

The human *UBE3A* gene encodes an E3 ubiquitin ligase that has three known protein isoforms^4,5^. Surprisingly little is known about the human isoforms; however, studies in which human isoforms were ectopically expressed in mouse cells indicate that the isoforms may have differences in localization and function^7,8^. Here we have used the CRISPR/Cas9 system to generate hESCs lacking these individual isoforms and studied their abundance and localization in both hESCs and neurons. We have also examined the phenotype of neurons lacking the individual isoforms. We have shown that human isoform 1 is the most abundant of the three isoforms, accounting for almost ninety percent of total UBE3A protein in these two cell types. We have also demonstrated that in both hESCs and neurons, the majority of UBE3A protein can be found in the cytoplasm, independent of which isoform is absent or present, although UBE3A is present in the nuclear and mitochondrial fractions, as well. Finally, we have shown that neurons lacking isoform 1, which accounts for approximately 84-88% of total UBE3A in both neurons and stem cells, display limited *in vitro* AS electrical phenotypes.

To examine the localization of the UBE3A isoforms, we used two approaches: subcellular fractionation and immunocytochemistry. Both approaches showed that isoform 1 is the predominant protein isoform in both hESCs and neurons. Loss of individual isoforms did not change the abundance or localization of remaining UBE3A, suggesting that each protein isoform may be translated from independent RNAs. Subcellular fractionation experiments revealed that the bulk of total UBE3A protein is found in the cytoplasmic portions of fractionated lysates and revealed that UBE3A is also located in the nuclear and mitochondrial fractions (which may also include some endoplasmic reticulum). Interestingly, immunocytochemistry experiments seemingly show that UBE3A is overwhelmingly localized to the nucleus of normal and isoform-null neurons. Rigorous experiments to confirm both fractionation and immunocytochemistry experiments with additional antibodies confirmed these findings. The distribution of UBE3A throughout normal neurons by the TEM suggests that the protein may be distributed somewhat evenly across all subcellular compartments. These seemingly disparate results may be explained by the relative proportion of cytoplasm versus nucleus in neurons. In most neurons, the nucleus occupies less than a tenth of the total cell volume^16^. We would then expect that an evenly distributed protein would be 10-fold more enriched in the cytoplasm versus the nucleus of neurons. Based on quantification of UBE3A and GAPDH protein levels in fractionated H9 neurons, we estimate that approximately 70% of UBE3A protein is found in the cytoplasm, leaving the remaining 30% to be localized to the nucleus and mitochondria, which accounts for ~10% of the cellular volume and may explain the apparent enrichment in the nucleus. Together, our data strongly suggest that in human neurons, UBE3A protein—including isoform 1—is found in all cellular compartments assayed.

Murine UBE3A has been shown to become increasingly nuclear during neuronal maturation as determined by immunocytochemistry.^9,17^ Furthermore, a recent publication reports that the short UBE3A isoform, which is most similar to human isoform 1, is predominantly nuclear.^9^ However, these studies only used immunocytochemistry to assess localization, and in many cases involved over-expressing UBE3A isoforms in wildtype mouse neurons, and thus may not accurately reflect the localization of endogenous isoforms. Fractionation data from our isoform 1 knockout neurons shows loss of cytoplasmic UBE3A, strongly suggesting that human isoform 1 is localized to both the nucleus and cytoplasmic compartments (**Figure 3c**). We cannot rule out the possibility that our hESC-derived neurons are too developmentally immature to display these changes in localization. However, a recent publication reporting the UBE3A distribution in human post-mortem brain sample shows both nuclear and cytoplasmic localization of UBE3A.^18^ Moreover, our isoform 1 null hESC-derived neurons appear to have UBE3A staining in the nucleus (**Figure 3g, Supplementary figures 4 and 6**), suggesting that the remaining two longer UBE3A isoforms are also localized in both cytoplasm and nucleus.

Isoform 1-null neurons lack approximately 84-88% of the total UBE3A protein and recapitulate some, but not all AS electrophysiological phenotypes. Isoform 1-null neurons have a more depolarized resting membrane potential (RMP) compared to controls or the other two isoform-null neuron (**Figure 4a**). It is possible that the difference in RMP between isoform 1-null neurons and their isogenic controls would be even more pronounced at an earlier age, such as 9 to 10 weeks *in vitro.* Although isoform 1-null neurons have a more depolarized RMP, nearly all cells fired single mature or mature trains of action potentials (**Figure 4b**). It is possible that a more subtly delayed development of action potential firing would be seen if we assayed this over a developmental time course in isoform 1-null neurons. It is interesting that we observe a milder physiological phenotype in neurons lacking isoform 1. This is consistent with the milder AS phenotype seen in patients with an analogous mutation in *UBE3A*^11^ Future studies are needed to examine the effects of the combined loss of two isoforms at a time to further elucidate their role in the electrical maturation of normal and AS neurons, as well as examine these phenotypes at various stages of neuronal development in our isoform-null neurons.

We have also shown here that isoform 1 accounts for the majority of UBE3A protein in hESCs and neurons, while isoforms 2 and 3 appear to account for very little protein at this stage. This is consistent with the relative abundances of long versus short isoforms of UBE3A in the mouse^9^. This knowledge of the relative abundance of the isoforms is important for the development of AS therapies. One promising therapeutic avenue currently being explored for AS is the introduction of a UBE3A transgene through vector-based therapies^10^, which typically can only contain one cDNA at a time. Replacement of all three human isoforms would likely require the development of three separate vectors, and is not feasible for current therapeutic approaches. Based on the results shown here, it is possible that delivery of isoform 1 alone may be a useful therapeutic approach, as it accounts for the majority of total UBE3A protein in human neurons *in vitro.* However, it is not known whether isoforms 2 and 3 have a specialized function not yet described, and it is possible that the different UBE3A isoforms have different ubiquitin ligase activities (i.e. more versus less active) or bind different co-factors. Future studies examining whether introduction of individual UBE3A protein isoforms into human AS iPSC-derived neurons is capable of restoring their phenotypes are needed to determine whether all isoforms are functionally interchangeable.

In summary, we have determined the relative abundance and localization of the three protein isoforms of UBE3A in human stem cells and neurons by genetic manipulation of the endogenous isoform translation start sites. We have shown that isoform 1 is the most abundant, and that UBE3A protein localizes to both the cytoplasm and nucleus in both cell types, independent of the specific isoform. We have also demonstrated that neurons lacking isoform 1 recapitulate some, but not all, AS physiological phenotypes. This knowledge is important for understanding the pathophysiology of AS cellular mechanisms and for the development of gene therapies to treat AS.

## Methods

### hESC culture and neural differentiation

hESCs were cultured on irradiated mouse embryonic fibroblasts and fed daily with hESC media (DMEM/F12 containing knockout serum replacement, L-glutamine + β-mercaptoethanol, non-essential amino acids, and basic fibroblast growth factor). hESCs were cultured in at 37°C in a humid incubator at 5% CO2. Cells were manually passaged every 5-7 days.

hESCs were differentiated into neurons using a modified version of the monolayer protocol. Neural induction was begun 2 days after passaging by culturing cells in N2B27 medium (Neurobasal medium, 1% N2, 2% B27, 2 mM L-glutamine, 0.5% penicillin/streptomycin, 1% insulin-transferrin-selenium). N2B27 medium was supplemented with fresh Noggin (500 ng/mL) for the first 10 days of differentiation. Neural rosettes were manually passaged onto poly-D-lysine and laminin coated plates using the Stem Pro EZ passage tool approximately 14 days after beginning neural induction. Neural progenitors were replated at a high density around 3 weeks of differentiation, switched to neural differentiation medium (NDM) around 4 weeks of differentiation, then plated sparsely for terminal differentiation at around 5 weeks. NDM consisted of neurobasal medium, 1% B27, 2 mM L-glutamine, 0.5% pen-strep, non-essential amino acids, 1 μM ascorbic acid, 200 μM cyclic AMP, 10 ng/mL brain-derived neurotrophic factor, and 10 ng/mL glial-derived neurotrophic factor. Neurons were maintained in culture until 10 to 12 weeks of differentiation, with the exception of the neurons in Figure 3f,g / Supplemental Figure 4, which were maintained until 15 weeks. The protocols for hESC maintenance and neuronal differentiation have been described previously^19,20^.

### CRISPR/Cas9-mediated genome editing

All CRISPRs were designed using MIT’s CRISPR design tool (http://crispr.mit.edu). sgRNAs were cloned into the pX459v2.0 vector (Addgene 62988), as previously described^21,22^ unless otherwise indicated. The 3’ sgRNA used to generate the UBE3A KO line was designed by the hESC/iPSC Targeting Core at the University of Connecticut Health Center and was cloned into the pX330 vector. The isoform 1 sgRNA has been published previously^15^. All sgRNA sequences used in this work are included in **Supplemental Table 1**. All ssODN sequences used to generate isoform KO lines and UBE3A HA-tagged lines are included in **Supplemental Table 2**.

Prior to electroporation or nucleofection, hESCs were treated with ROCK inhibitor (Y-27632; Selleck Chemicals) for 24 hours. hESCs were then singlized using Accutase (Millipore) and electroporated using the Gene Pulser X Cell (BioRad) or nucleofected using the Amaxa 4D Nucleofector (Lonza). For generation of the Isoform1 and UBE3A KO lines, hESCs were electroporated in PBS with 10 μg of CRISPR plasmid and, for the Isoform1 KO line, 8 μl of single-stranded oligonucleotide (ssODN) template (100 μM). For generation of the Isoform2 KO and Isoform3 KO lines, and the HA-tagged UBE3A lines, hESCs were nucleofected with 2 μg of CRISPR and 3 μl ssODN (100 μm) using the P3 Primary Cell Kit L (Lonza). hESCs were then plated onto puromycin-resistant (DR4) irrMEFs at low density, supplemented with ROCK inhibitor and L755507 (5 μM, Xcessbio), which has been shown to improve efficiency of homology directed repair^23^. 24 hours after plating, cells underwent selection for 48 hours with puromycin (0.5 – 1 μg/ ml). Puromycin resistant colonies were screened by conventional PCR (UBE3A KO and UBE3A HA-tagged lines) or conventional PCR followed by restriction digest (Isoform KO lines) 11-14 days after plating. Putative clones were plated onto regular irrMEFs and successful genome editing was confirmed by Sanger sequencing. Primer sequences used to generate the hESC lines in this paper are listed in **Supplemental Table 3**.

### Subcellular Fractionation

Subcellular fractionation was performed using the Cell Fractionation Kit – Standard (Abcam) according to the manufacturer’s instructions with the following modifications: 1) Protease Inhibitor Cocktail III (EMD Millipore) was added to 1x Buffer A at the beginning of the fractionation protocol at a 1:1000 dilution; 2) a whole cell lysate was collected following the first lysis step in Buffer B by collecting 1/6 of the volume of lysate, and downstream volumes used in the protocol were adjusted accordingly; and 3) whole cell and nuclear fractions were sonicated at the end of the protocol to sheer DNA using the following settings: [3 seconds on, 3 seconds off] x 2 at 30% amplitude. All fractions were stored at −80°C until use.

### Western Blot

Equal volumes of lysate from each fraction were separated by SDS-PAGE using 4-20% TGX Stain-Free mini gels (BioRad). Protein was transferred to PVDF membrane using the TransBlot Turbo system (BioRad). Membranes were blocked in TBS Odyssey Blocking Buffer (LI-COR, Inc.) for 1 hour at room temperature then incubated in blocking buffer containing primary antibodies overnight at 4°C. Membranes were washed with TBS-T (Tris-buffered saline plus 0.1% Tween-20) at room temperature, incubated in blocking buffer containing IRDye Secondary Antibodies (LI-COR, Inc.) for 1 hour at room temperature, then washed again in TBS-T. All washes were done three times for 10-12 minutes each at room temperature. Membranes were imaged using the Odyssey imaging machine and software (LI-COR, Inc.). Images were quantified using Image Studio Lite (LI-COR, Inc.). The following primary antibodies were used: rabbit anti-UBE3A (1:3000; Bethyl A300-351), rabbit anti-UBE3A (1:3000; Bethyl A300-352), mouse anti-UBE3A (1:1000; Sigma E8655), rabbit anti-LMNB1 (1:5000; Abcam), mouse anti-GAPDH (1:10000; Millipore); rabbit anti-BAK (1:1000; Cell Signaling Technologies). The following secondary antibodies were used at a concentration of 1:10000: IRDye 800 CW Donkey anti-Rabbit, IRDye 680RD Goat anti-Rabbit, and IRDye 680RD Goat anti-Mouse (LI-COR, Inc.). A Western blot containing only nuclear fractions from all 4 hESC and neuron lines was performed with the following changes from above: blocking and antibody incubations were done in 5% BSA, anti-rabbit HRP-conjugated secondary antibody (1:3000; Cell Signaling Technologies) was used, and the blot was imaged using Clarity Western ECL substrate (BioRad) on the ChemiDoc Touch imaging system (BioRad).

### Immunocytochemistry

hESCs and neurons were grown on glass chamber slides (Thermofisher Scientific) for immunocytochemistry. Cells were fixed using room temperature 4% paraformaldehyde for 10 min then permeabilized using 0.5% PBS-Triton X 100 (PBS-T) for 10 min at room temperature. Cells were blocked in 0.1% PBS-T containing 2% bovine serum albumin and 5% normal goat serum. Cells were incubated in primary antibody in blocking buffer overnight at room temperature then washed with PBS. Cells were then incubated in secondary antibody in blocking buffer for 3 hours at room temperature then washed with PBS. Cells were mounted with ProLong Gold Anti-Fade Hard Set with DAPI (ThermoFisher Scientific) and allowed to set overnight at room temperature before imaging. The following primary antibodies were used: mouse anti-UBE3A (1:500; Sigma SAB1404508 clone 3E5), mouse anti-UBE3A (1:500; Sigma E8655 clone E6AP-330), rabbit anti-OCT4 (1:200; Stemgent 09-0023), chicken anti-MAP2 (1:10000; Abcam ab3592), rabbit anti-NeuN (1:300; Abcam ab177487). The following secondary antibodies were used: anti-mouse Alexa Fluor 488 (1:400; Invitrogen), anti-rabbit Alexa Fluor 594 (1:400; Invitrogen), anti-chicken Alexa Fluor 647 (1:250, abcam). Confocal images were acquired with a 63x oil immersion objective (NA 1.4) on a Zeiss 780 confocal system mounted on an inverted Axio Observer Z1 (Carl Zeiss, Germany). Images were averaged twice, with pixel size of 0.26 μm, pixel dwell time of 1.27 seconds, and scan time of 2.34 seconds. Three randomly chosen areas of the chamber slide were images for each cell line. Z-series were done with a 0.2 μm interval and maximum intensity projection images were created using Fiji (ImageJ) software.^24^

### Electrophysiology

Neurons were plated onto glass coverslips around 5 weeks of differentiation. Whole-cell voltage and current clamp recordings were performed at 12 weeks of differentiation as previously described^15^.

### qRT-PCR

cDNA was synthesized using the High-Capacity cDNA Reverse Transcription Kit (Thermo Fisher Scientific) according to manufacturer’s instructions. Quantitative RT-PCR was performed using Taqman Gene Expression Assays and Mastermix (Thermo Fisher Scientific) on the Step One Plus (Thermo Fisher Scientific). Reactions were performed in technical duplicates, with GAPDH Endogenous Control Taqman Assay used as the housekeeping gene for normalization. Gene expression was quantified using the ΔΔC_t_ method.

### Immunogold Transmission Electron Microscopy

Neurons were plated for terminal differentiation onto plastic 4 well plates at 6 weeks of differentiation. At 10 weeks, neurons were fixed using 4% paraformaldehyde at room temperature for 30 min. Cells were washed in PBS at room temperature, then blocked and permeabilized by incubating in 0.1% PBS-T containing 2% bovine serum albumin and 5% normal goat serum for 30 min at room temperature. Cells were incubated overnight at room temperature in blocking buffer containing primary antibody (mouse anti-UBE3A (Sigma SAB1404508 clone 3E5)) at 1:250). Cells were washed at room temperature in PBS then incubated in secondary antibody for 1 hour at room temperature. Secondary antibody (goat antimouse Nanogold Fab) was diluted 1:200 in 1% BSA. Cells were washed several times in PBS then washed in deionized water. Gold enhancement was performed for 4 min using GoldEnhance (Nanoprobes). Cells were rinsed in deionized water to stop enhancement reaction.

To prepare cells for electron microscopy, cells were rinsed for 5 min in 0.1 M Cacodylate buffer, then post-fixed in 1% OsO_4_, 0.8% Potassium Ferricyanide in 0.1 M Cacodylate buffer for 30 min at room temperature. Cells were then washed 5 times in deionized water for 5 min per wash, block stained in 1% Uranyl Acetate in deionized water for 30 min at room temperature, then washed again 3 times (5 min each) in deionized water. Next cells were dehydrated in ethanol (5 min in 50% EtOH, 5 min in 75% EtOH, 5 min in 95% EtOH, then 3 incubations in 100% EtOH for 5 min each), then infiltrated in 100% resin overnight at room temperature. The following day, fresh 100% resin was added and the cells were polymerized at 60°C for 48 hours. Thin sections of 70-80 nm were cut on the Ultramicrotome Leica EM UC7 and sections placed on Cu grids. Sections were counterstained with 6% Uranyl Acetate in 50% methanol for 4 min. Images were acquired with the Hitachi H-7650 Transmission Electron Microscope. Electron microscopy preparation and imaging was performed by the UConn Health Central Electron Microscopy Facility.

### Statistics

Statistical analysis was performed used SPSS software. Univariate ANOVA was performed to compare levels UBE3A protein levels in Western blots (Fig 2b-e, Fig 3b-e), and to compare resting membrane potential (Fig 4a) and spontaneous synaptic activity (Fig 4c,d) between normal and isoform null cells. Chi square tests were performed to compare action potential firing patterns (Fig 4b) between normal and isoform null cells.

## Acknowledgements

The authors would like to thank members of the Chamberlain and Levine labs for helpful discussions during the course of this project. This work was funded by NIH grants R21HD091823-02 and R01HD094953-01 and an Angelman Syndrome Foundation grant to SJC, an Angelman Syndrome Foundation fellowship to NDG, and a Connecticut DPH Regenerative Medicine Research Fund grant 14-SCDIS-UCHC-01 to SJC, ESL, and LML.

## Author Contributions

CLS and SJC conceived of and designed the study, collected and analyzed data, and wrote the manuscript. SK generated the UBE3A-HA tagged lines and performed characterization of said lines. JEB performed all confocal microscopy except for Supplemental Figure 7, which was performed by NDG. JJF performed whole-cell electrical recordings and the data was analyzed by DG. ESL provided critical revision of the manuscript.

