## Supplemental Data for "Abundance and localization of human UBE3A protein isoforms"

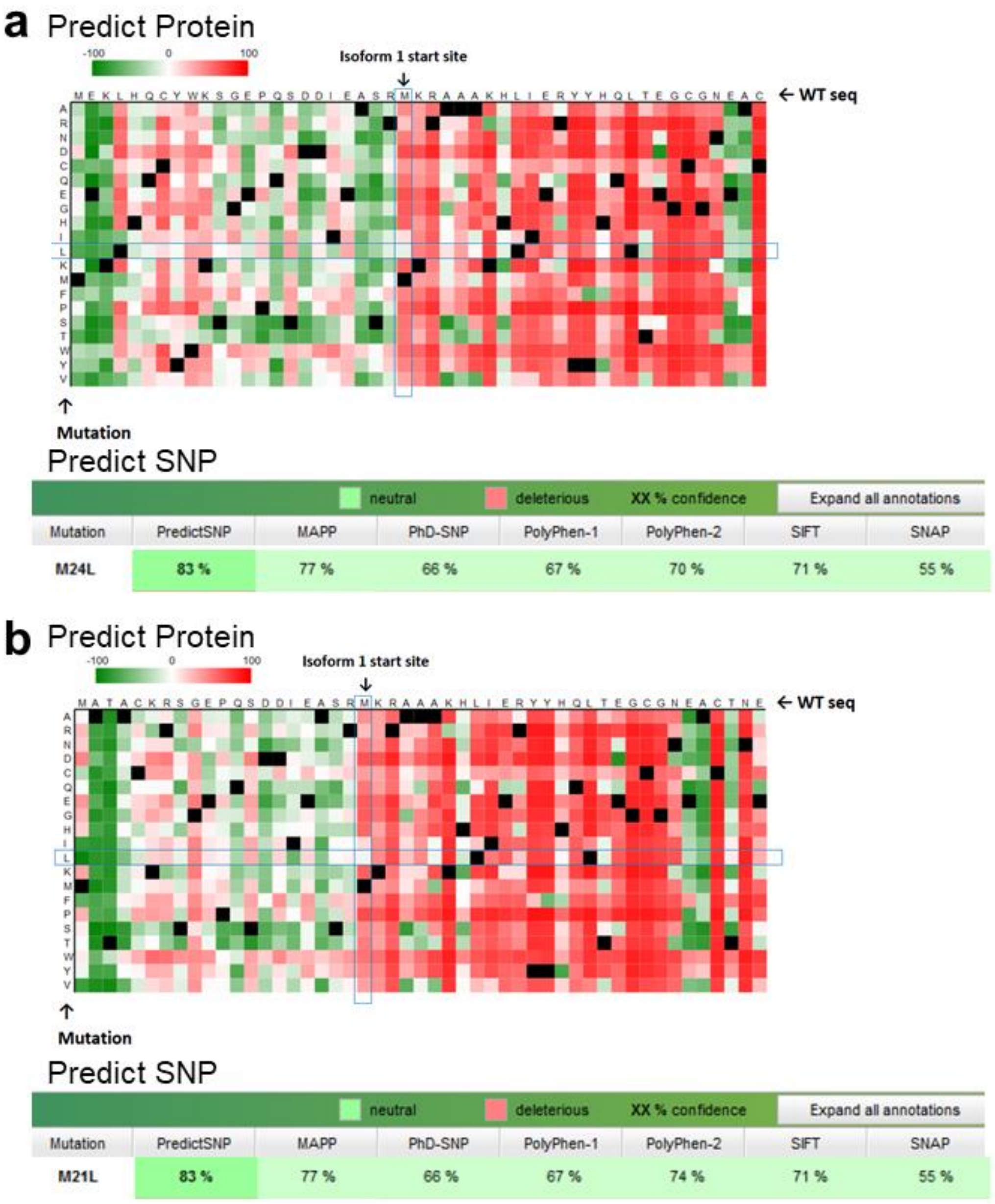


**Supplemental Figure 1. Effect of proposed amino acid change on isoform 2 (a) and isoform 3 (b) protein.**

Top: results from Predict Protein software showing effects of every possible amino acid substitution of the methionine (blue box). Bottom: results from Predict SNP software showing predicted effects of L to M substitution from 7 different structure prediction algorithms.

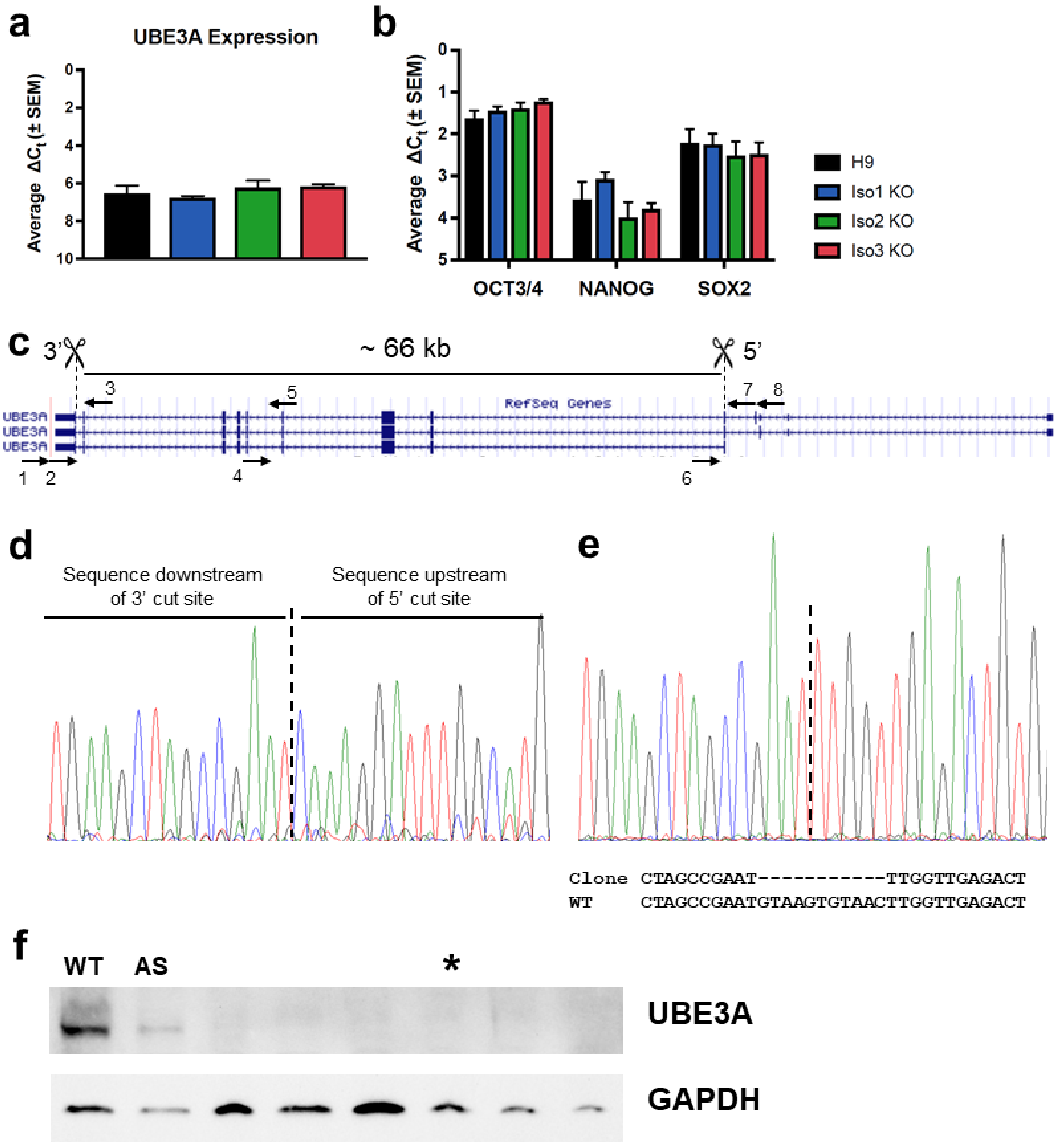


**Supplemental Figure 2. Characterization of isoform-null hESCs and generation of UBE3A KO hESCs**

**a** qRT-PCR results showing UBE3A expression in H9 and isoform-null hESCs **b** qRT-PCR results showing expression of pluripotency genes in H9 and isoform-null hESCs **c** Schematic showing location of two CRISPRs used to generate UBE3A KO line. Arrows indicate primers used for screening clones. **d** Sanger sequencing of PCR product spanning the deletion (primer 1 + primer 8) showing continuous sequence from upstream of 5' cut site to downstream of 3' cut site. **e** Sanger sequencing of the non-deleted allele showing an 11 bp deletion at the 5' cut site. **f** Western blot showing knockout of UBE3A in hESCs from UBE3A KO clone (*). WT = normal iPSCs, AS = Angelman Syndrome iPSCs. Other lanes in the image are other clones that were screened following genome editing. Error bars in **a**,**b**: standard error of the mean.

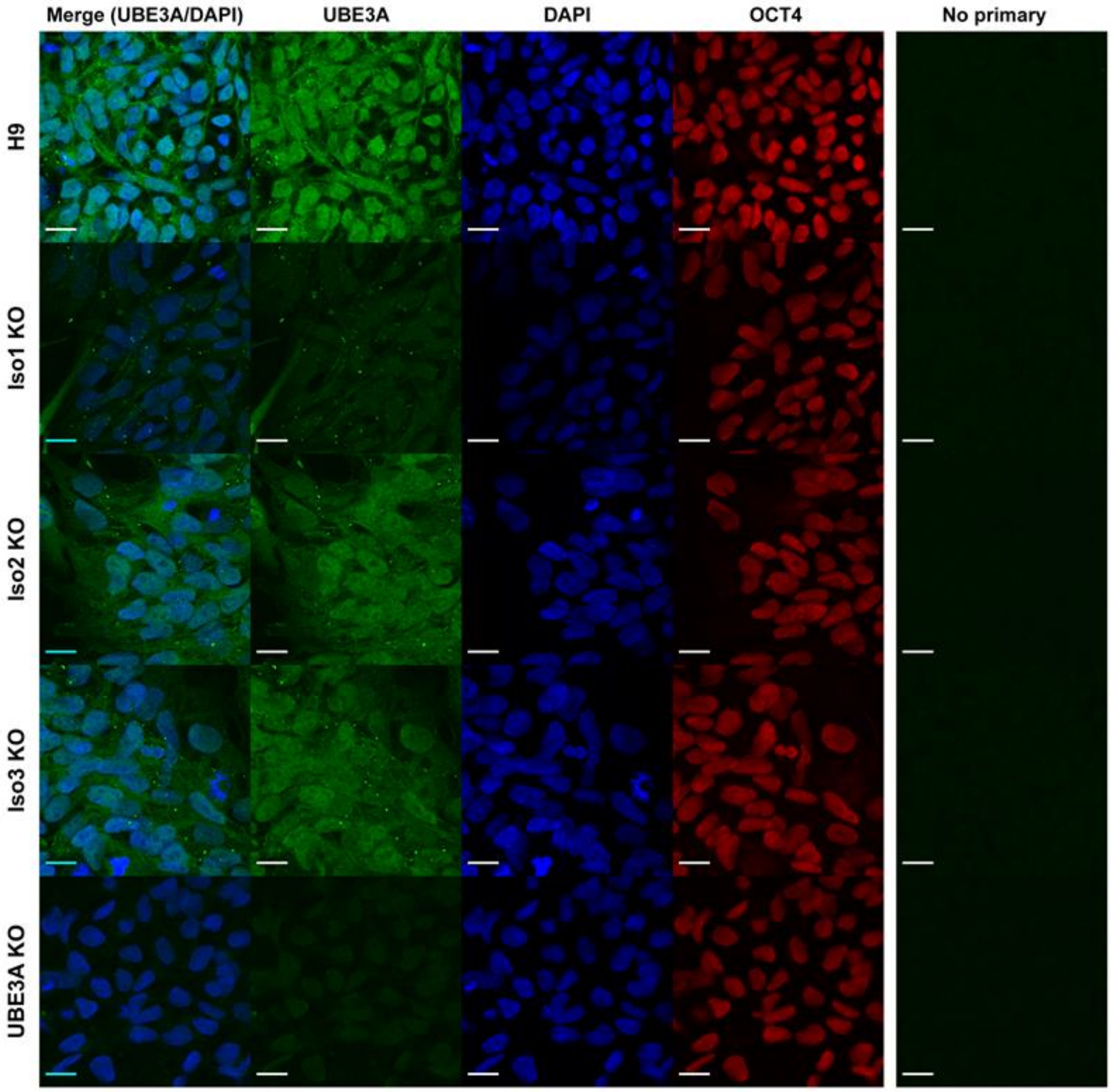


**Supplemental Figure 3. UBE3A expression in isoform-null hESCs by immunocytochemistry**

Abundant expression of UBE3A is seen in the cytoplasm in all isoform-null cell lines, indicating that the localization of UBE3A does not differ upon loss of any isoform. Isoform 1-null hESCs show dramatically reduced protein levels. UBE3A KO hESCs were included as a negative control. OCT4 staining was included to show localization of a known nuclear protein. Scale bar 20 μm.

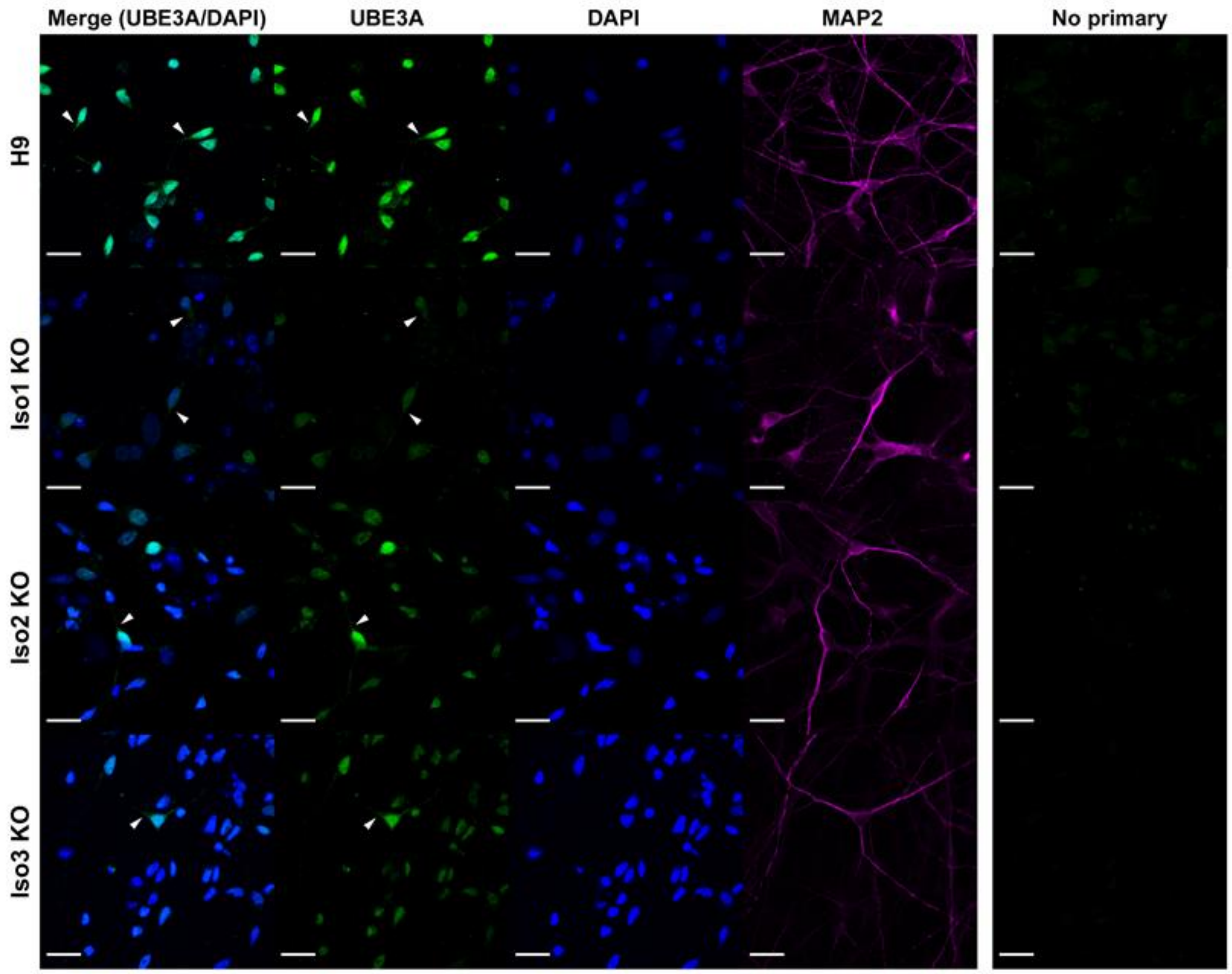


**Supplemental Figure 4. UBE3A expression in neurons by immunocytochemistry**

UBE3A signal appears to localize predominantly to the nucleus by immunostaining in neurons, in contradiction to fractionation results. Scale bar 20 μm. Arrowheads indicate areas in the neuron where UBE3A is expressed outside of the nucleus.

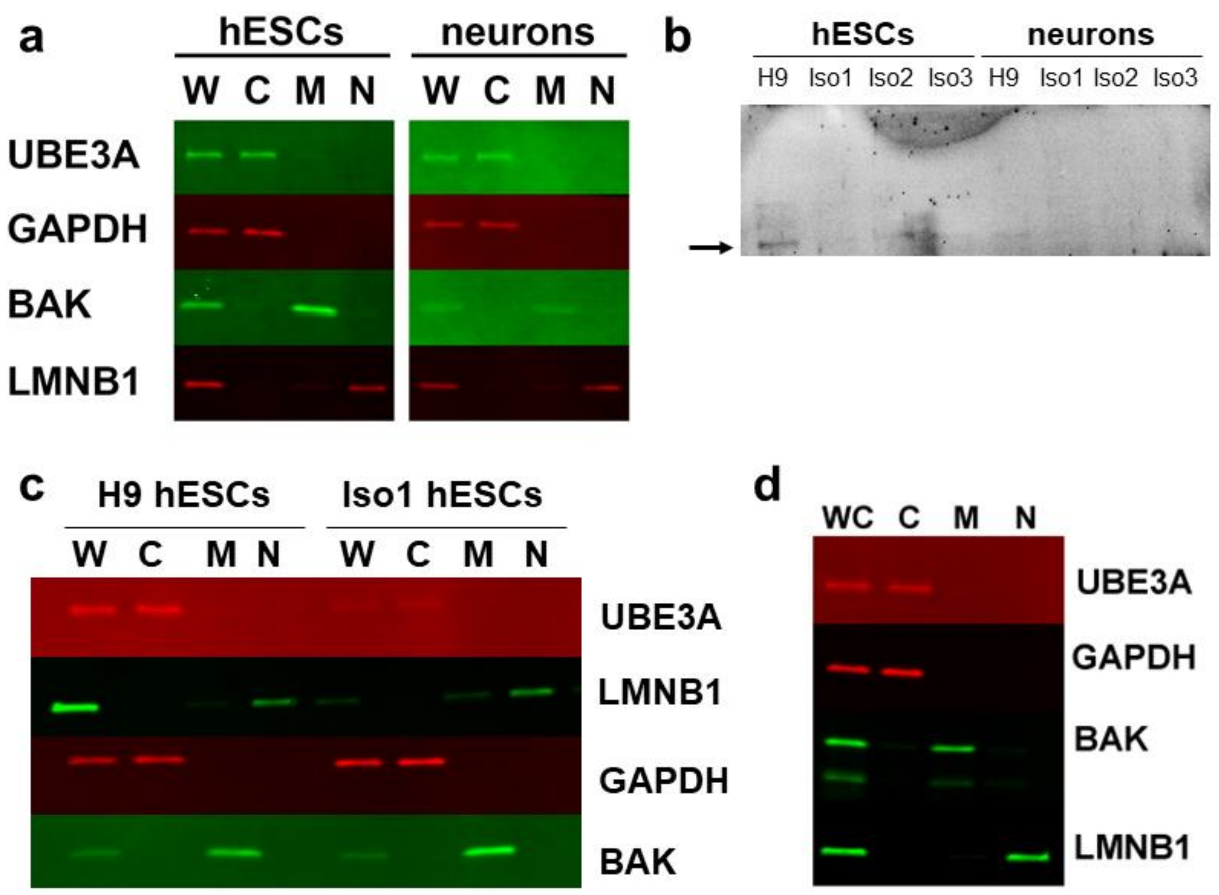


**Supplemental Figure 5. Confirmation of fractionation results**

**a** Confirmation of fractionation results in H9 hESCs and neurons using a second UBE3A primary antibody **b** Overexposure of nuclear fractions from wildtype and isoform-null hESCs and neurons show faint nuclear UBE3A signal **c&d** Western blot using primary antibody used for ICC on lysates from wildtype and isoform 1-null hESCs (**c**) and wildtype neurons (**d**) indicates that fractionation results are consistent using multiple antibodies. W = whole cell fraction; C = cytoplasmic fraction; M = mitochondrial fraction; N = nuclear fraction

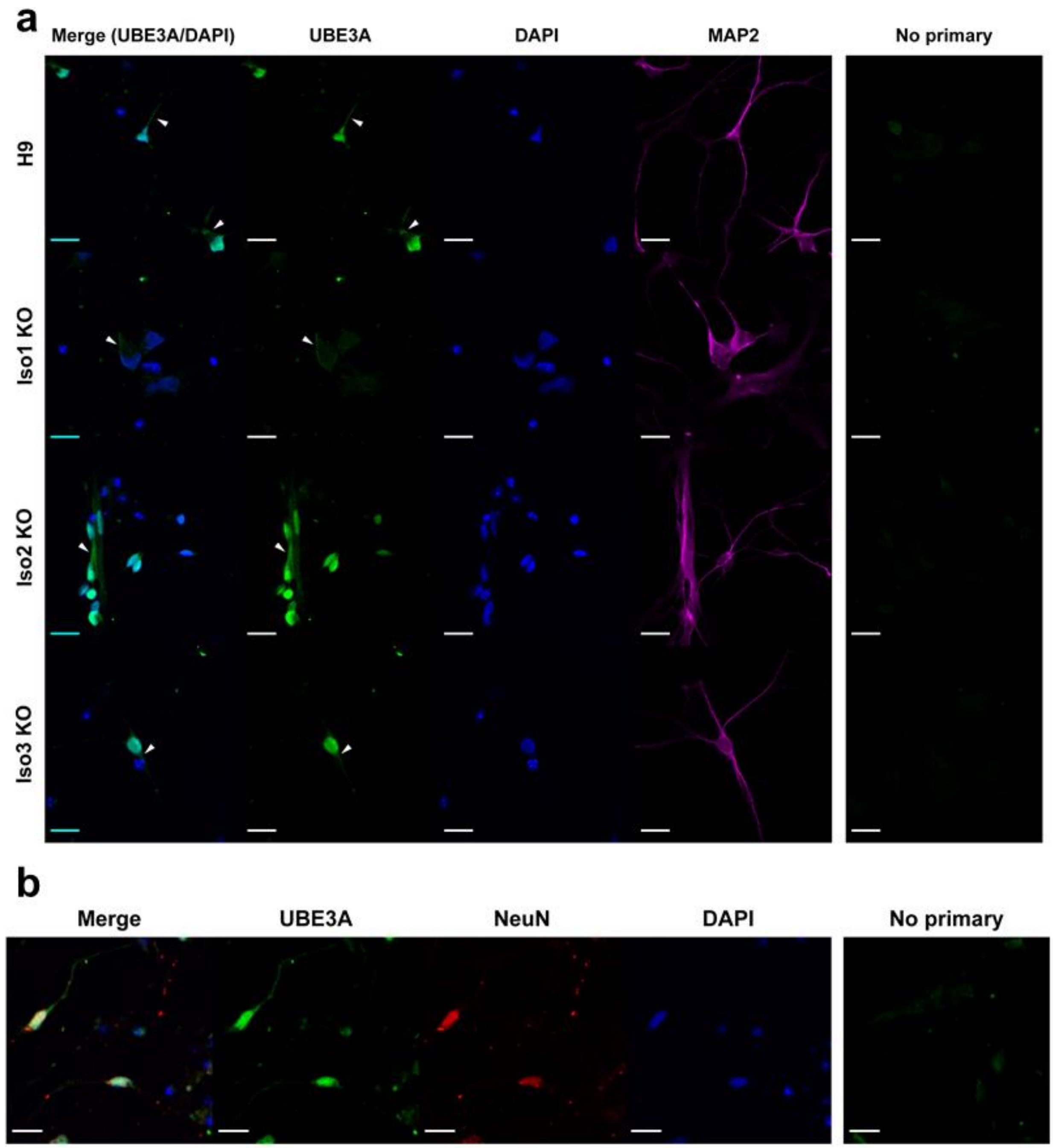


**Supplemental Figure 6. Confirmation of immunocytochemistry results using a second primary antibody**

**a** Using a second antibody against UBE3A shows similar patterns of expression of UBE3A in hESC-derived neurons. **b** UBE3A partially colocalizes with NeuN, but not completely. Scale bar 20 µm. Arrowheads indicate areas in the neuron where UBE3A is expressed outside of the nucleus.

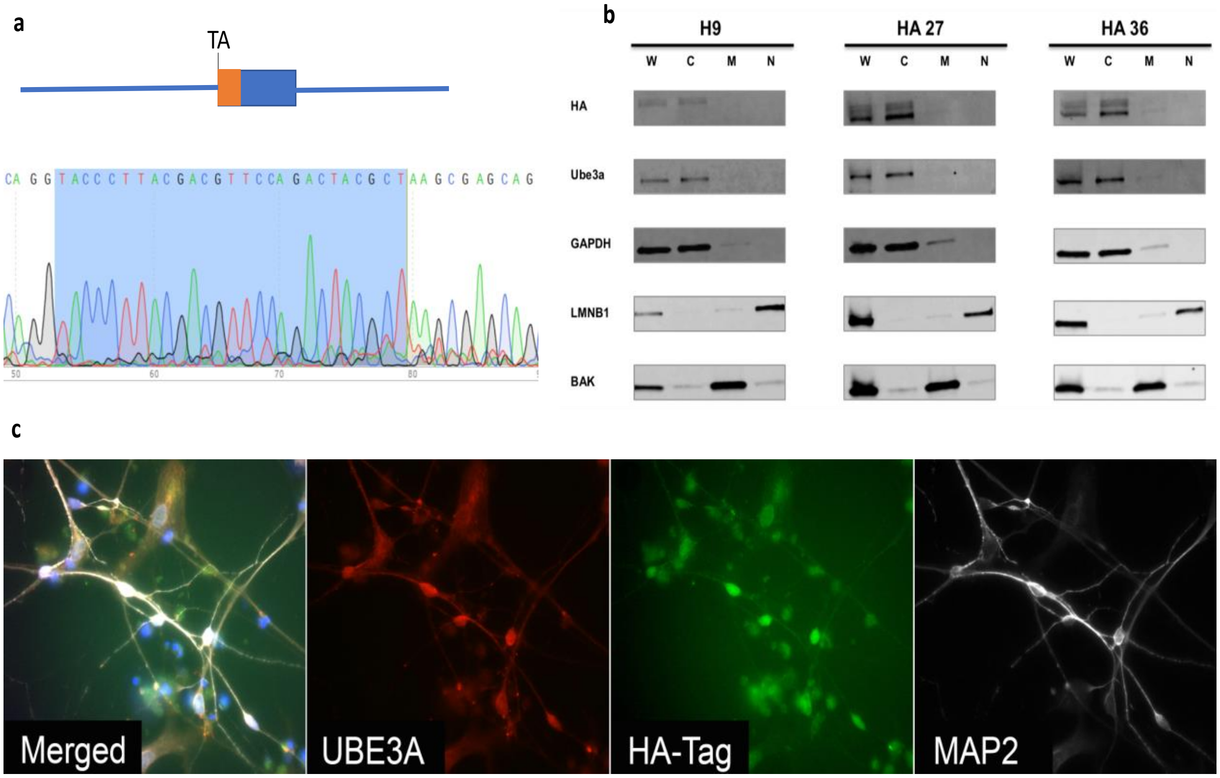


**Supplemental Figure 7. HA-tagged UBE3A confirms localization of UBE3A**

**a** Strategy for HA-tag insertion into isoform 1 of UBE3A. Direction of transcription is right to left, and AT from translational start site of UBE3A isoform 1 is indicated. Sanger sequence of HA-tag from HA-27 hESC line is shown with HA sequence highlighted. **b** Fractionation of two independent hESC lines harboring homozygous insertion of the HA-tag into UBE3A isoform 1. **c** Immunocytochemistry of HA-27 hESC-derived neurons stained with antibodies against UBE3A (red), HA (green), and MAP2 (far-red, shown in black and white).

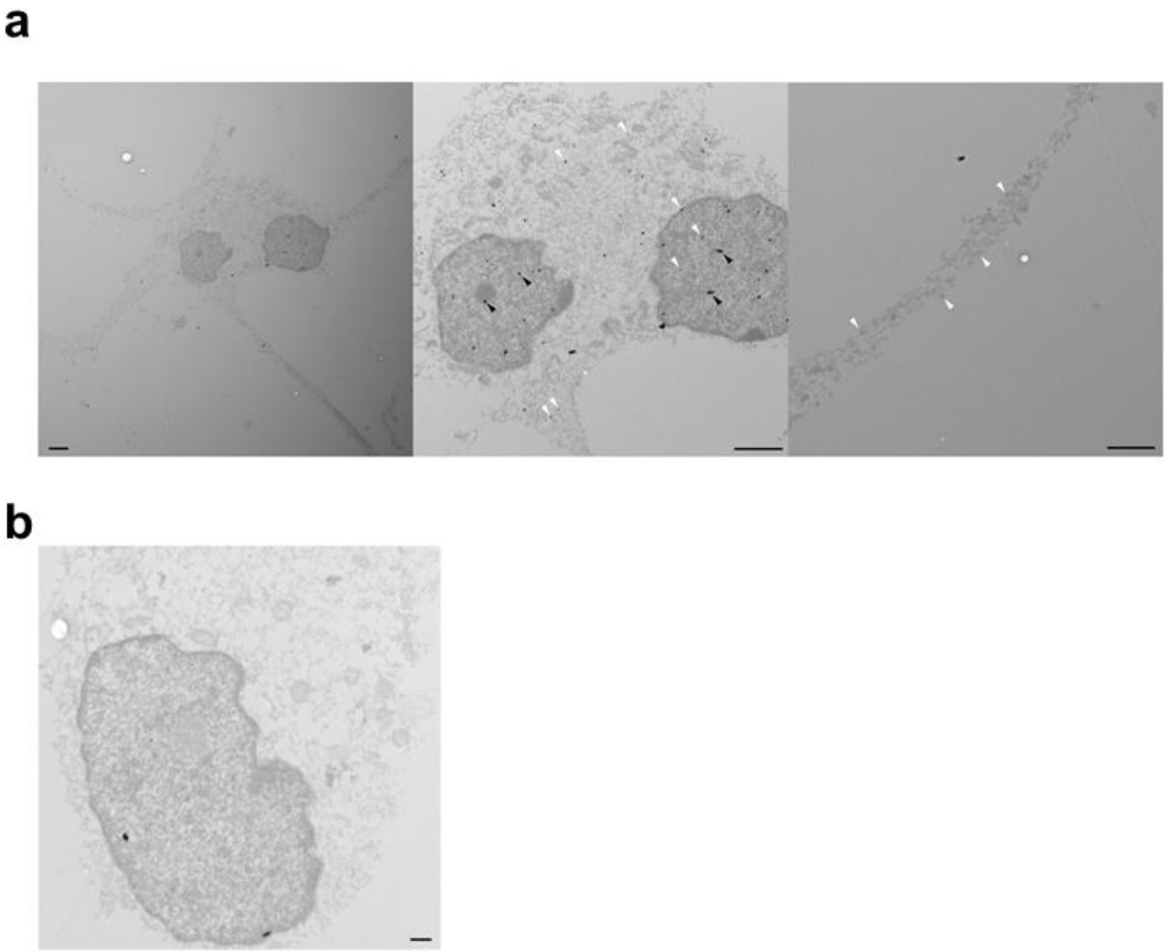


**Supplemental Figure 8. Transmission electron microscopy images show diffuse UBE3A localization in cytoplasm and nucleus in H9 neurons**

**a** Immunogold labeling shows diffuse UBE3A staining (white arrowheads) in the nucleus and soma (middle image) as well as in the neuronal processes (right image). Examples of non-specific puncta are indicated by black arrowheads. Scale bar 2 μm **b** No primary control image. Scale bar 500 nm.

| Supplemental Table 1. sgRNAs used to generate stem cell lines | |
|---|---|
| Cell Line | sgRNA sequence + **PAM** |
| Isoform 1 start site mutation / UBE3A KO (5') | **CCG**AATGTAAGTGTAACTTGGTT |
| Isoform 2 start site mutation | AAAAGGAGTGGCTTGCAGGA**TGG** |
| Isoform 3 start site mutation | ATCACCCTGATGTCACCGAA**TGG** |
| UBE3A KO (3') | AGGCCATCACGTATGCCAA**AGG** |
| UBE3A-HATag | **CCA**CCAGTTAACTGAGGGCTGTG |

Supplemental Table 1. Sequences of sgRNAs used for genome editing of hESCs to generate isoform-null and UBE3A KO lines.

| Supplemental Table 2. ssODNs used to generate isoform-null stem cell lines | |
|---|---|
| Isoform 1 start site mutation | AGAACCTCAGTCTGACGACATTGAAGCTAGTCGATTGTAA<br>GTGTAACTTGGTTGAGACTGTGGTTCTTAT |
| Isoform 2 start site mutation | ATTCAAATGGTGGCTCACTTCCAATAACACTGGTGAAGCTTCTCGA<br>GCCTGCAAGCCACTCCTTTTACCTCCACTGTAACTCTCTAGGAGAG |
| Isoform 3 start site mutation | TGTAAAATAATTCAAAATTACCTTTTACAAGCTGTTGCAAGTCGGTG<br>ACATCAGGGTGATCACAGCTTTGAGTCACTGATTAAAAA |
| HA Tag insertion at Isoform 1 start site | AGCTTTGTACTTGAGAGCTGTACTAATCACTGTGCTTATTGTTTGAA<br>TGTTTGGTACAGGTACCCTTACGACGTTCCAGACTACGCTAAGCGA<br>GCAGCTGCAAAGCATCTAATAGAACGCTACTATCACCAGTTAACTG<br>AGGGCTGTGGAAATGAAGCCTGCACGAATGAGTTTTGT |

Supplemental Table 2. Sequences of single-stranded oligonucleotides (ssODNs) used as templates for homology-directed repair during CRISPR genome editing.

| Supplemental Table 3. Primer Sequences Used for Genome Editing | | |
|---|---|---|
| Label within Figures | Primer Name | Sequence |
| L / 6 | Iso1R | CTGCTACCAGGGAAGCAAAA |
| M | Iso1SeqF | TTCTTTCATGTTGACATCTTTAATTTT |
| N / 7 | Iso1F | GCTTATAATGGCTTGTCTGTTGG |
| O | Iso2R | TGAAACAATAACCAAATAACATTGG |
| P | Iso2SeqR | TCTTGATTTGAATCGCAGAAAA |
| Q | Iso2F | TCAGTAGCCACTATCAAAGACCT |
| R | Iso3R | TTTTTGAACAATGAATTGGGTTT |
| S | Iso3SeqF | AGCCTACGCTCAGATCAAGG |
| T | Iso3F | TTTTTGAACAATGAATTGGGTTT |
| 1 | UBE3A del R2 | TGGGACACTATCACCACCAA |
| 2 | UBE3A del R1 | CCCACATGTCCCCAATAAAG |
| 3 | 3' cut site upstream F | GGCAACTTGGTAGTTACACAACA |
| 4 | Internal R | GCACTTGAGAAAACAATGTCCA |
| 5 | Internal F | TGAAACACTTTGGAAATGTAGCC |
| 8 | UBE3A del F1 | AGTCCAACCCTTAAAATAAATGTG |

Supplemental Table 3. Primer sequences used for PCR screening of clones for isoform start site genome editing. Primers labeled with letters refer to **Figure 1a**, while primers labeled with numbers refer to **Supplemental Figure 2c**
